# Zoom in on Ca^2+^ pattern and ion flux dynamics to decode spatial and temporal regulation of cotton fiber growth

**DOI:** 10.1101/2021.02.08.430284

**Authors:** Jia-Shuo Yang, Jayakumar Bose, Sergey Shabala, Yong-Ling Ruan

## Abstract

Cotton fibers are single-celled trichomes initiated from ovule epidermis prior to anthesis. Thereafter, the fibers undergo rapid elongation for 20 d before switching to intensive cell wall cellulose synthesis. The final length attained determines fiber yield and quality. As such, cotton fiber represents an excellent single cell model to study regulation of cell growth and differentiation, with significant agronomical implications. One major unresolved question is whether fiber elongation follows a diffusive or a tip growth pattern. We addressed this issue by using cell biology and electrophysiological approaches. Confocal imaging of Ca^2+^ binding dye, fluo-3 acetoxymethyl (Fluo-3), and *in situ* microelectrode ion flux measurement revealed that cytosolic Ca^2+^ was evenly distributed along the elongating fiber cells with Ca^2+^ and H^+^ fluxes oscillating from apical to basal regions of the elongating fibers. These findings demonstrate that, contrary to growing pollen tubes or root hairs, cotton fiber growth follows a diffusive, but not the tip growth, pattern. Further analyses showed that the elongating fibers exhibited substantial net H^+^ efflux, indicating a strong activity of the plasma membrane H^+^-ATPase required for energy dependent solute uptake. Interestingly, the growing cotton fibers were responding to H_2_O_2_ treatment, know to promote fiber elongation, by a massive increase in the net Ca^2+^ and H^+^ efflux in both tip and basal zones, while non-growing cells lacked this ability. These observations suggest that desensitization of the cell and a loss of its ability to respond to H_2_O_2_ may be causally related to the termination of the cotton fiber elongation.

**One sentence summary:** Confocal imaging of Ca^2+^ patterning and *in situ* microelectrode ion flux measurements demonstrate that, contrary to growing pollen tubes or root hairs, cotton fiber growth follows a diffusive, but not the tip growth, pattern.

## Introduction

Cotton (*Gossypium*) fibers are single-celled trichomes initiated from a quarter to one third of ovule epidermal cells about 16 to 24 h prior to anthesis (Ruan et al., 2000; Wang et al 2021). Following their protrusion from the ovule epidermis, the fiber cells undergo rapid cell elongation for about 20 d before switching to intensive secondary cell wall cellulose synthesis that lasts for 15 to 20 d (Ruan, 2005; Ruan, 2007). By maturity, the fiber cells can reach ~3 and ~5 cm long, in the cultivated species of *G. hirsutum* and *G. barbadense*, respectively, rendering cotton fiber one of the longest cells in the plant kingdom (Kim and Triplett, 2001; Ruan et al., 2001). As such, cotton fiber represents an attractive single-cell model to study cell expansion (Ruan et al., 2001; Ruan et al., 2004; Shan et al., 2014; Yu et al., 2019), cellulose synthesis (Amor et al., 1995; Ruan et al., 2003) and cell patterning (MacHado et al., 2009; Walford et al., 2011; Tian et al., 2020; Wang et al., 2020). Apart from its significance in studying fundamental plant biology, cotton fiber is the most important source of cellulose for the global textile industry. Therefore, advance in fiber biology will help to design innovative approaches to improve cotton yield and quality using advanced breeding or gene technology.

As outlined above, one extraordinary feature of the cotton fiber cells is its fast and sustained elongation at the average rate of 1500 to 2500 μm per day for about 20 d, with little or no increase in cell diameter that remains at around 10 to 15 μm (Ruan, 2007). Given this remarkable cellular characteristic and the fact that the fiber length is a key determinant of cotton yield and quality, there has been intensive research on the regulation of cotton fiber elongation over the last three decades. To this end, early studies established that the microtubules in elongating fiber cells are arranged transversely along the longitudinal axis (Seagull, 1990), which guided cellulose deposition accordingly, thereby forcing fibers to elongate unidirectionally (Ruan, 2007), a phenomenon confirmed recently using transgenic approach (Yu et al., 2019). On the other hand, cell biology and gene expression analyses revealed a temporary closure of plasmodesmata (PD), due to the deposition of callose at the fiber base, which coincided with the maximal expression of the plasma membrane sucrose transporter and expansin genes at the onset of rapid elongation from 10 d post anthesis, DPA (Ruan et al., 2001; Ruan et al., 2004). These findings underpin a model of the fiber elongation driven by the cell turgor and relaxed wall expansibility (Ruan, 2007). A follow-up study revealed that the fiber PD gating is under a tight control of the sterol-mediated callose degradation, and an increase in the duration of PD closure via silencing a sterol carrier gene activated the expression of genes encoding both H^+^-coupled sucrose transporters and -uncoupled clade III SWEETs (Zhang et al., 2017b). Recent molecular genetic studies and genome sequencing have identified a number of regulatory genes and networks underlying cotton fiber elongation (e.g. Shan et al., 2014; Hu et al., 2019).

Despite of the aforementioned progress, major questions remain regarding the spatial and temporal regulaiton of cotton fiber elongation. A key unresolved issue is whether the fiber cells elongate evenly along the the whole growh axis, or possess the tip-based growth pattern as that found in morphologically-similar structures such as root hairs or growing pollen tubes (Hepler et al., 2013; Mangano et al., 2018; Nakamura and Grebe, 2018). In other words, does cotton fiber elogation follow a diffusive pattern or a tip growth model? A related question is whether and how the growth pattern may change during elongation. In this context, cotton fiber cells are known lacking zonation of the endoplasmic reticulum, Golgi bodies and mitochondria or viscles in the tip region (Tiwari and Wilkins, 1995) that are characteristic of the tip-based growth (Yang, 1998). This is indicative of no polarized deposition of the new cell wall material for the apex expansion. By imaging fibers expressing fluorescent-tagged cytoskeleton proteins, Yu et al. (2019) observed that microtubles are organzed transversely during fiber elongation, instead of being in bundles in parallel to the growth axis as typically observed in tip growth cells such as pollen tubes (Yang, 1998), a finding consistent with an early report (Seagull, 1992). These results support a diffusive growth model of the cotton fiber proposed previously (Ruan, 2007).

However, counter arguments against the diffusive growth model arise from several Ca^2+^ imaging studies, where elevated levels of Ca^2+^ were observed at the tip region of 2-5 DPA fibers (Huang et al., 2008; Zhang et al., 2017a), which is consistent with a tip growth pattern observed in elongating pollen tubes (Chen et al., 2009; Qu et al., 2012; Gu et al., 2015; Suwińska et al., 2017) and root hairs (Monshausen et al., 2008; Fan et al., 2011). A close examination of those reports, however, revealed that the fluorescent Ca^2+^ signal was observed in a much extensive region of the cotton fiber tips, often spanning 100 to 500 μm from the edge of the tip (e.g. Huang et al., 2008), which does not match the geometric distribution of Ca^2+^ in either pollen tubes or root hairs that employ tip-based growth mechanism. Here, the hallmark of the tip growth is a localized elevation in the cytosolic free Ca^2+^ that is confined to only 5~ 25 μm region from the tip boundary (e.g. Fan et al., 2011; Gu et al., 2015; Gilroy et al., 2016). This, together with a lack of proper positive and negative controls and resolutions required to differentiate cytosol from other subcellular compartments such as vacuole in the previous reports on cotton fibers (Huang et al., 2008; Zhang et al., 2017a), raises a question of the validity of those findings on Ca^2+^ localization and the conclusions drawn (e.g. Qin and Zhu 2011). Also, reliance of fluorescence dyes alone come with a caveat of the possible methodological issues of its loading in various types of cells and/or regions that could be related to the properties of the cell walls. This calls for a need to employ some other techniques to provide an unequivocal answer to the above question.

Physiologically, intracellular Ca^2+^ concentration is determined by Ca^2+^ influx and efflux across cellular membranes, with tip-based growing cells characterized with pronounced Ca^2+^ oscillation in the tip, but not in the shank region (Gilroy et al., 2016). Similar oscillatory patterns have been reported for pollen tubes (Holdaway-Clarke et al., 1998; Damineli et al., 2017) that also employ tip-based growth mechanism. There have been no studies thus far on the spatial behaviour of Ca^2+^ oscillation along the longitudinal axis of the growing cotton fibers.

To determine whether the cotton fiber cell follows a tip or diffusive growth pattern and to better understand the spatial and temporal regulation of fiber elongation, we have combined two advanced complementary techniques, namely fluorescent imaging using Ca^2+^ binding dye, fluo-3 acetoxymethyl (Fluo-3 thereafter) (Zhang et al., 1998; Zhang et al., 2015) and non-invasive ion-selective microelectrode technique, MIFE (Microelectrode Ion Flux Estimation, Shabala et al., 1997; Wu et al., 2020). Our analyses revealed that cytosolic Ca^2+^ was evenly distributed along the elongating fiber cells, with Ca^2+^ and H^+^ oscillations occurring in both apical and basal regions of the elongating fibers at 5 and 10 d post anthesis (DPA), a phenomenon absent in elongated fibers at 20 DPA. These findings demonstrate that cotton fiber growth clearly follows the diffusive but not tip-based growth pattern. Pharmacological experiments using various Ca^2+^ channel blockers on cultured cotton ovules indicate the involvement of both voltage-dependent and -independent Ca^2+^ channels in the cotton fiber elongation. Our analyses further uncovered several novel patterns including findings that (i) the elongating, but not elongated, cotton fibers exhibited massive H^+^ efflux with strongest observed at 10 DPA, the onset of rapid fiber elongation, indicating strong plasma membrane H^+^-ATPase activity; (ii) H_2_O_2_ stimulated Ca^2+^ and H^+^ efflux from elongating, but not elongated, fibers and (iii) H_2_O_2_ application promoted K^+^ uptake into and efflux from growing and non-growing fibers, respectively. This data is discussed in the context of regulation of cell expansion by cellular energy status, ion homeostasis and H_2_O_2_-mediated signal transduction.

## Results

### Validation on visualization of Ca^2+^ in the cytosol of cotton fiber cells

To visualize Ca^2+^ signals in growing cotton fiber cells, cotton seeds with fibers attached were harvested on day of anthesis (0 DPA) and pre-loaded with ester form of Fluo-3 to produce the intracellular Ca^2+^-binding fluorescent probe, Fluo-3, following the cleavage of the ester group by the cytosolic esterase (Zhang et al., 1998; Zhang et al., 2015). A set of experiments was firstly conducted to ascertain that the Fluo-3 fluorescent signals were emitted from the binding of cytosolic Ca^2+^, but not from that of other subcellular compartments.

Confocal imaging of the free-hand sections counterstained with Calcofluor White for cellulose revealed green fluorescent signals of the Ca^2+^-binding Fluo-3 in the region between cell walls and vacuoles (Fig. 1, A and B). Here, the Calcofluor White-labeled cell walls exhibited blue fluorescence only, whereas the vacuoles (asterisks in Fig. 1, A) showed no fluorescence, flanked by the Fluo-3 green fluorescence in the middle. The observation indicates that no Fluo-3 was produced in the cell wall and vacuolar compartments in our procedure, which could happen due to the presence of esterase in the extracellular matrix or leakage of tonoplasts (Zhang et al., 1998). To determine if the green fluorescence of Fluo-3 was sensitive to changes in Ca^2+^ level, we cultured 0-d cotton seed for 5 d on the same but Ca^2+^ free BT medium. This intervention abolished the green fluorescence, indicating no or little Ca^2+^ left in the fiber cells following the Ca^2+^ starvation treatment, although some residual signals remained visible in the underneath seed coat (Fig. 1, C). Elongating pollen tubes and root hairs are known to have high Ca^2+^ level in the cytosol of their respective apical regions, thus representing ideal positive controls. Loading both cell systems with Fluo-3 resulted in a strong fluorescent signal in their tip regions (Fig. 1, D and E), as expected (e.g. Fan et al., 2011; Suwińska et al., 2017).

**Figure 1.**
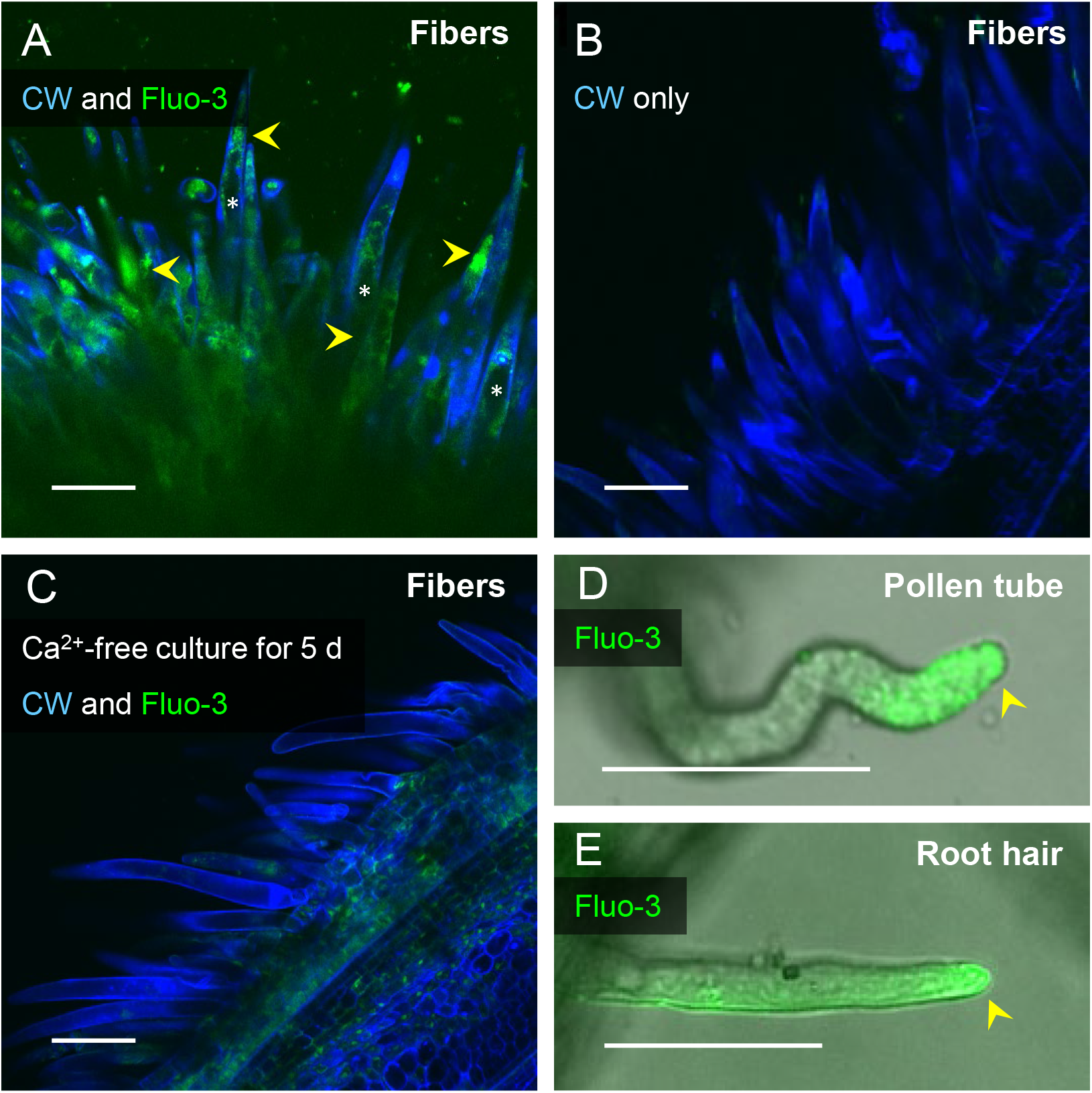
Validation of Ca2+ indicator Fluo-3 in cotton fibers, pollen tubes and root hairs. Green fluorescence came from the membrane-impermeable intracellular Ca^2+^ indicator Fluo-3, whereas blue fluorescence reflected the cell wall indicator Calcofluor White (CW) binding to cellulose. Cotton seeds (5 days post anthesis, DPA) with fibers attached were co-stained with Fluo-3 and CW (A), or with CW only (B). (C) Cotton fibers derived from seeds cultured in a Ca^2+^ -free BT medium were co-stained with Fluo-3 and CW at 5 DPA. Note, the presence of Ca^2+^ fluorescence signals (arrowheads) in (A) outside the vacuoles (asterisks) but its absence in the negative controls of (B) and (C). (D and E) Representative images of cotton pollen tube and cotton root hair, respectively, stained with Fluo-3, showing Ca^2+^ fluorescence at the tip regions. Scale bars in (A, B and C) = 100 μm, in (D and E) = 50 μm.

To provide an unequivocal evidence that the fluorescence signal emitted from the Ca^2+^-binding Fluo-3 did come from the cytosol of the elongating fibers, plasmolysis was performed on cultured seed and fiber with 0.5 or 1.0 M sorbitol. In comparison with the non-plasmolysed control of 5-d cotton fibers, which exhibited a green fluorescence of Fluo-3 inside the cell wall but outside the vacuole (Fig. 2, A), plasmolysis induced by 0.5 M sorbitol caused the fiber protoplast being pulled away from the cell wall, leaving a gap between the cell wall displaying a blue fluorescence signal from Calcofluor White and the narrow strip exhibiting green fluorescence (Fig. 2, B and C). A triple labelling with Fluo-3, Calcofluor White and RH-414, a plasma membrane-specific staining (Zhang et al., 2015) revealed that the Fluo-3 green fluorescence resided within the red fluorescence of RH-414 for plasma membrane (Fig. 2, C). In comparison to that treated with 0.5 M sorbitol (Fig. 2, B and C), the effect of plasmolysis became more evident following incubation with 1.0 M sorbitol, in which the region emitting Ca^2+^-bound Fluo-3 green fluorescence was pulled further away from the cell wall with the vacuole becoming highly shrunken (Fig. 2, D) due to osmotically driven efflux of water (Ruan et al., 2000). These data established clearly that the Ca^2+^-bound Fluo-3 green fluorescence was emitted from the cytosol of the elongating cotton fiber cells.

**Figure 2.**
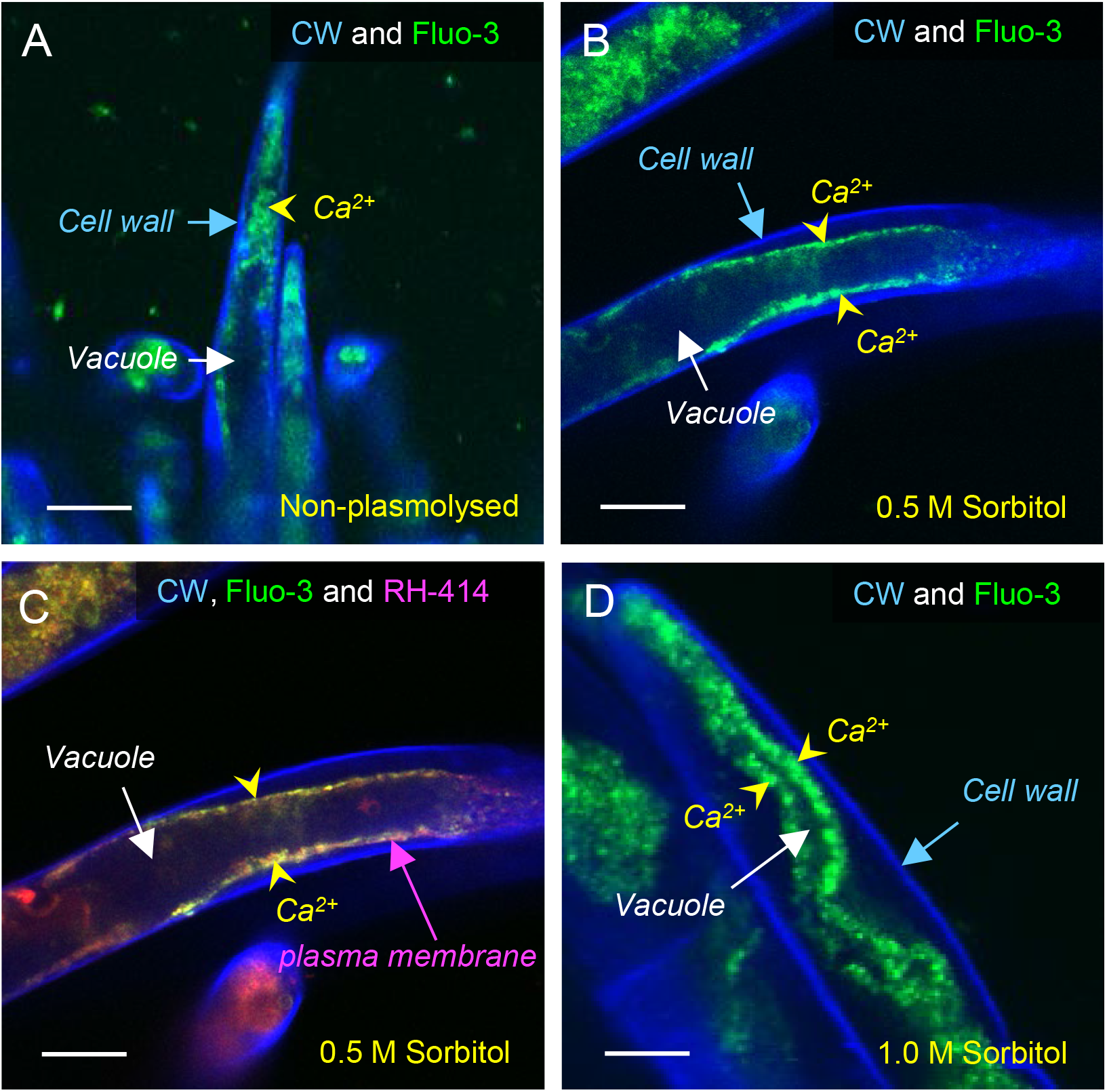
Localization of Ca^2+^ in the cytosol of elongating fiber cells. Ca^2+^ was indicated by green fluorescence emitted from Fluo-3. Cell wall and plasma membrane were labelled with Calcofluor White (CW, blue fluorescence) and with RH-414 (red fluorescence), respectively. 5 d cotton seeds with fibers attached were incubated for 30 min in PBS medium (A), or the PBS with sorbitol at 0.5 M (B and C) or 1 M (D) before confocal imaging. Note, in comparison with the control (non-plasmolysed in (A), plasmolysis resulted in protoplast being pulled away from the cell wall (B, C), a phenomenon becoming more evident under 1.0 M sorbitol treatment (D). In all cases, Ca^2+^ was localized in the cytosol between the cell wall and vacuole. Scale bars = 15 μm.

### Cotton fiber cells exhibited even distribution of the cytosolic Ca^2+^ from tip to base throughout the elongation period

Having validated the procedure of using Fluo-3 to visualize cytosolic Ca^2+^ in fiber cells as illustrated above, we systemically imaged Ca^2+^ localization patterns in fibers across the entire elongation phase from 0 to 20 DPA as well as the peak stage of the secondary cell wall cellulose synthesis at 30 DPA. Representative images from selected stages were shown in Figure 3.

**Figure 3.**
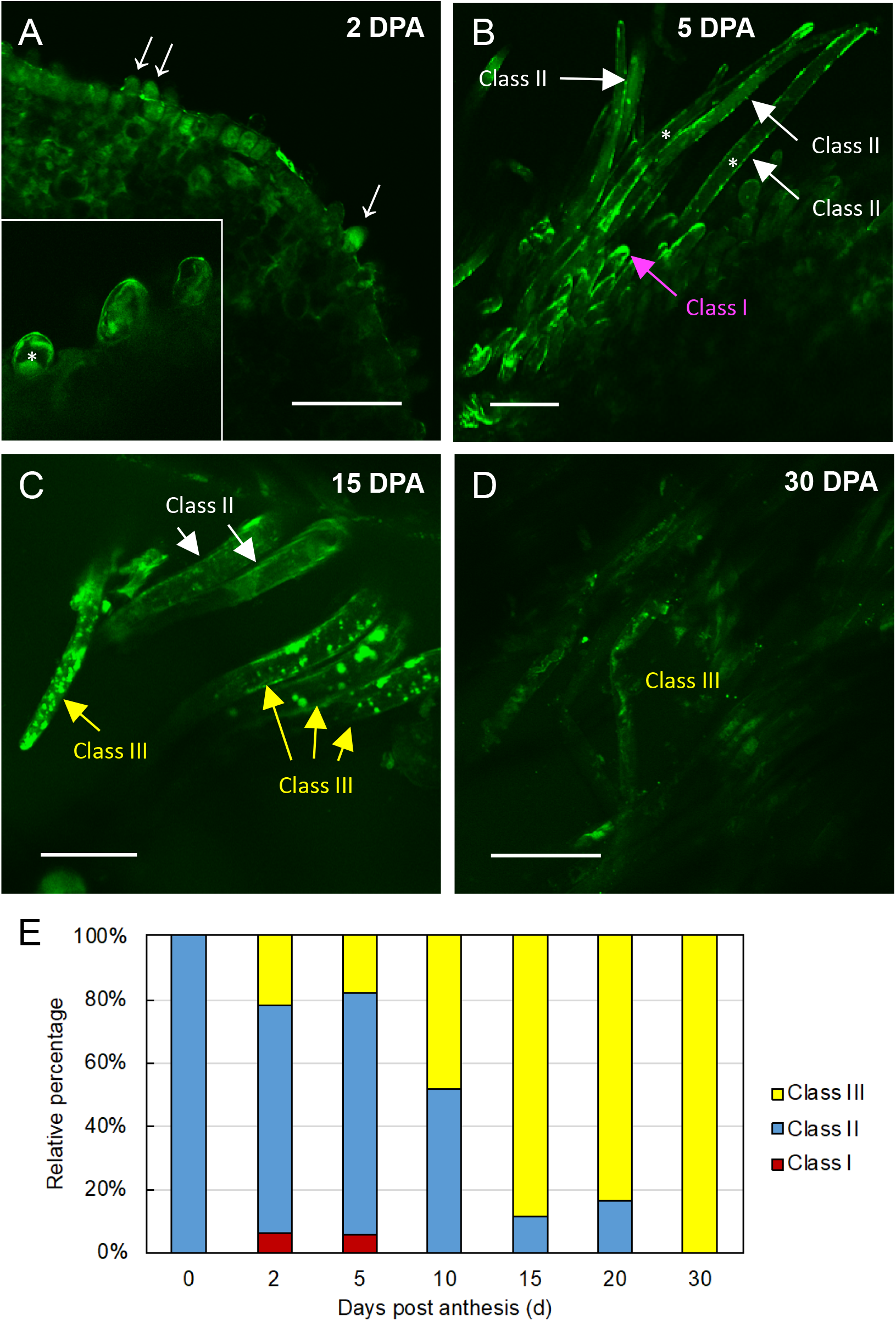
Dynamics of Ca^2+^ patterning in cotton fibers from elongation to rapid cell wall cellulose synthesis. Ca^2+^ was indicated by green fluorescence emitted from Fluo-3 labelling. (A) Young fiber cells elongated from seed epidermis (arrows) at 2 DPA, showing Ca^2+^ signals in the cytosol (insert). (B) Elongating fiber cells at 5 DPA, exhibiting Ca^2+^ signals occasionally concentrated to the tip area but mostly in the peripheral regions of the fibers, termed as pattern class I and II, respectively. Asterisks in (A and B) indicate vacuoles. (C) Cotton fibers at 15 DPA, a transition phase from elongation to secondary cell wall cellulose synthesis. The fluorescent Ca^2+^ signals became aggregated or patchy, categorized as pattern class III. (D) Cotton fibers at 30 DPA, undergoing intensive cellulose synthesis, displayed much reduced Ca^2+^ signals, in comparison with that in the early stages (A, B, and C). Scale bars = 100 μm. I The relative percentages of classes I, II and III Ca^2+^ patterns across fiber development, which were calculated by counting at least 30 individual fiber cells randomly selected from 10 seeds at each stage.

The Ca^2+^ fluorescent signals were observed throughout the cytosols of almost all of the 0~2 d fiber initials (Fig. 3, A and insert). As the fiber cells elongate, the even distribution of cytosolic Ca^2+^ became more evident, reflected by the narrow strip of the fluorescent signals from tip to base with the non-fluorescent vacuole sitting in the centre (Fig. 3, B). Depending on the angle of the confocal imaging, some fiber cells exhibited tip-like fluorescent pattern at the first glance. However, a close examination revealed most of them displayed the fluorescence in the basal region as well (Class II in Fig. 3, B). Fiber cells with a true tip-localization of Ca^2+^ (Class I) was only sporadically observed. The uniform distribution of cytosolic Ca^2+^ pattern persisted to the end of elongation at 20 DPA. However, from 10 DPA onwards, an increased proportion of the fiber cells showed patchy fluorescence for Ca^2+^, a phenomenon highlighted in 15 DPA fibers where boundaries between vacuole and cytosol became less defined and patchy signals (Class III) were seen across the whole protoplast area (Fig. 3, C). By 30 DPA, the fluorescence became very faint and patchy (Fig. 3, D).

The relative percentages of fibers exhibiting classes I, II and III Ca^2+^ patterns at each stage is shown in Figure 3 (E). As one can see, 0-d fiber initials were all in the class II category, exhibiting uniform Ca^2+^signals throughout the cytosol. Tip-based location of the fluorescent Ca^2+^ signals was observed only in less than 5% of young fibers at 2-5 DPA and none in any other stages examined (Fig. 3, E). Interestingly, patchy pattern of Ca^2+^ signals (class III) increased from about 20% of 2-5 d fibers to ~ 48% at 10 DPA and then to ~ 80% at 15 to 20 DPA when elongation slows down and terminates (Ruan, 2007). By 30 DPA, all fibers displayed a patchy pattern of Ca^2+^, although the fluorescent signals became very weak at this stage (Fig. 3, D and E).

### Steady net Ca^2+^ fluxes were similar between tip and basal regions of fiber cells

Net Ca^2+^ fluxes were measured from apical and basal parts of the cotton fiber cells (illustrated in Supplemental Figure 1) using MIFE technique. No significant (at *P* < 0.05) difference in the steady-state Ca^2+^ flux was found between the two regions, with net Ca^2+^ uptake of 25 to 40 nmol m^−2^ s^−1^ measured (Fig. 4, A). These steady-state Ca^2+^ fluxes were largely independent of the cell age and not statistically different (*P* < 0.05) between growing (5- and 10-d old) and non-growing (20 d old) fibers (Fig. 4, A). Also similar and not statistically different (except one value for 20 d old base) were steady net K^+^ fluxes (Fig. 4, C). At the same time, net H^+^ fluxes showed a clear age-dependent trend (Fig. 4, B), showing a maximal net H^+^ efflux at 10 d. These results are consistent with the previous observations that 10 d is the turning point of switching from symplastic to apoplastic pathway when cells enter the rapid phase of elongation and probably possess strongest ATPase activity (Ruan et al., 2001). In non-growing 20-d fibers, this H^+^ efflux is ceased, with net H^+^ uptake detected (Fig. 4, B). At any age, the magnitude of net H^+^ fluxes was higher in the tip compared with the base of the fiber (Fig. 4, B) suggesting higher metabolic activity in this region.

**Figure 4.**
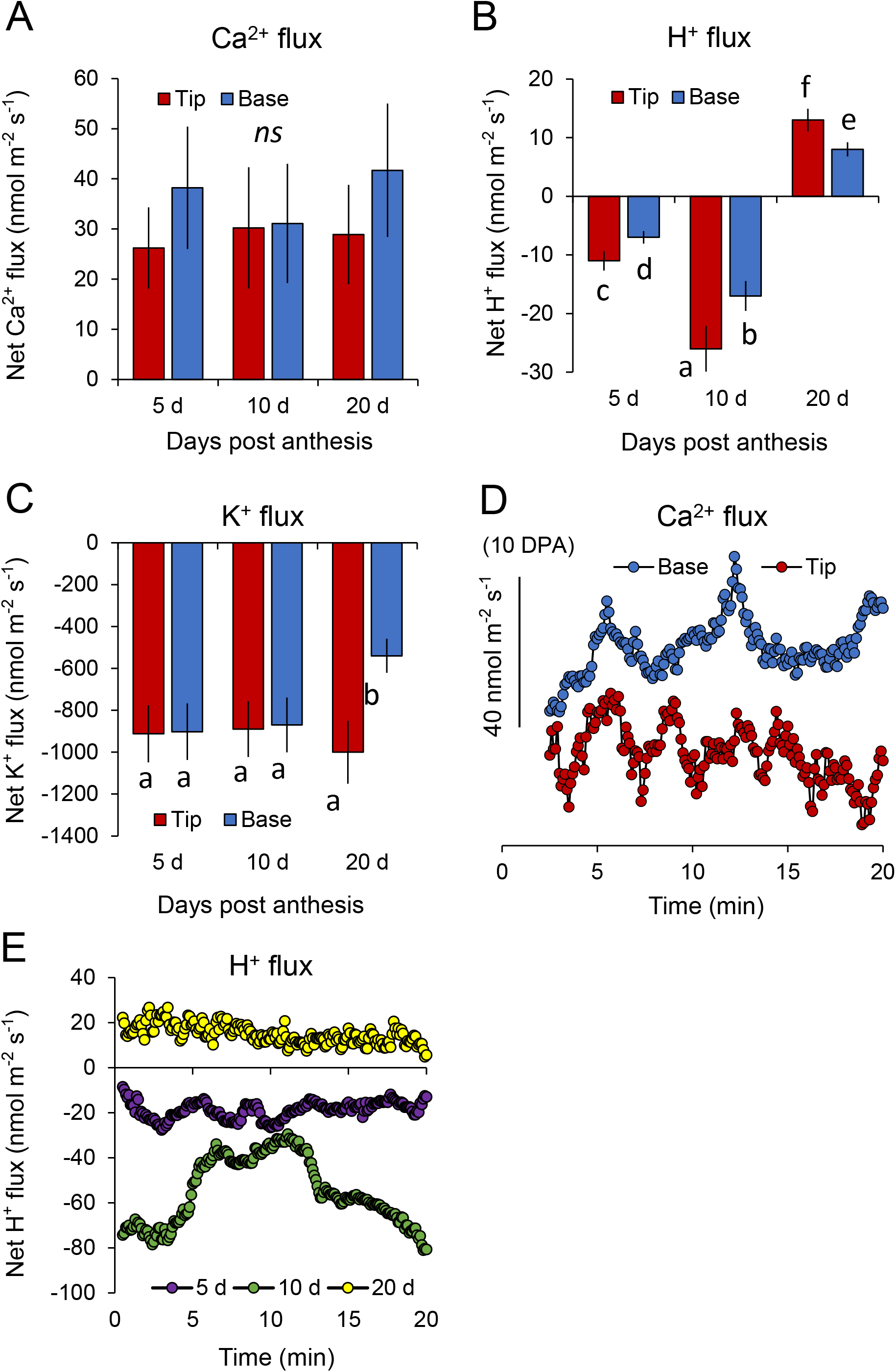
Net ion flux profiles across the plasma membrane of the cotton fiber cells measured by the non-invasive microelectrode MIFE technique. Panel A to C show mean values of steady-state net Ca^2+^ (A), H^+^ (B), and K^+^ (C) fluxes measured from the tip and basal regions of growing (5 and 10 DPA) and non-growing (20 DPA) cotton fiber cell. Mean ± SE (n = 5 to 8). Data labelled with different low-case letters is significant at *P* < 0.05. *ns* = not significant. (D) oscillations in net Ca^2+^ measured from the tip and basal regions of growing cotton fiber (10 DPA). (E) dynamics of net H^+^ fluxes measured from the tip region of the cotton fiber cells as a function of its age. The ultradian H^+^ oscillations were observed from growing (5 and 10 DPA) but not in non-growing (20 DPA) cells. One (of 6) typical example is shown for each treatment. The sign convention for all MIFE data is “influx positive”.

### Elongating fibers displayed ion flux oscillations in both tip and basal regions

One of the hallmarks of growing pollen tubes and root hairs is a presence of clearly pronounced ultradian (a minutes’ range of periods) oscillations that are observed in the cell tip but absent in the basal region (Monshausen et al., 2008; Zhou et al., 2014; Mangano et al., 2018; Hoffmann et al., 2020). Here, net Ca^2+^ ion flux oscillations were observed in both tip and base regions of the 10-d cotton fiber (Fig. 4, D), further supporting the concept of the diffusive fiber growth. The frequency of these oscillations were similar to those reported in the literature for root hairs and pollen tubes and ranged between 2 and 8 min in period. These oscillations were absent in non-growing (20 d old) fibers (as illustrated in Fig. 4, E for H^+^ flux data). Given a well-established role of Ca^2+^ oscillations in a broad range of plant developmental and adaptive responses (Tian et al., 2020), it is, therefore, plausible to suggest that the difference between growing and non-growing fiber cells may be potentially related to their ability to encode some vital information by means of Ca^2+^ flux oscillations.

### H_2_O_2_ stimulate Ca^2+^ and H^+^ efflux from elongating but not elongated fiber cells

Previous studies on both root hairs (Foreman et al., 2003) and pollen tubes (Lee and Yang, 2008) have revealed an important role of H_2_O_2_ as a component of the cell growth mechanism. H_2_O_2_ was also shown to be involved in cotton fiber growth (Tang et al., 2014). Maintaining cytosolic ROS homeostasis has been implicated in regulating cotton fiber elongation, with low H_2_O_2_ levels likely promoting elongation (Li et al., 2007). To check how ROS may impact on ion flux dynamics, we studied net ion flux responses to physiologically relevant concentration of H_2_O_2_ in two zones of growing (10 d) and 20-d non-growing fiber cells (Fig. 5).

**Figure 5.**
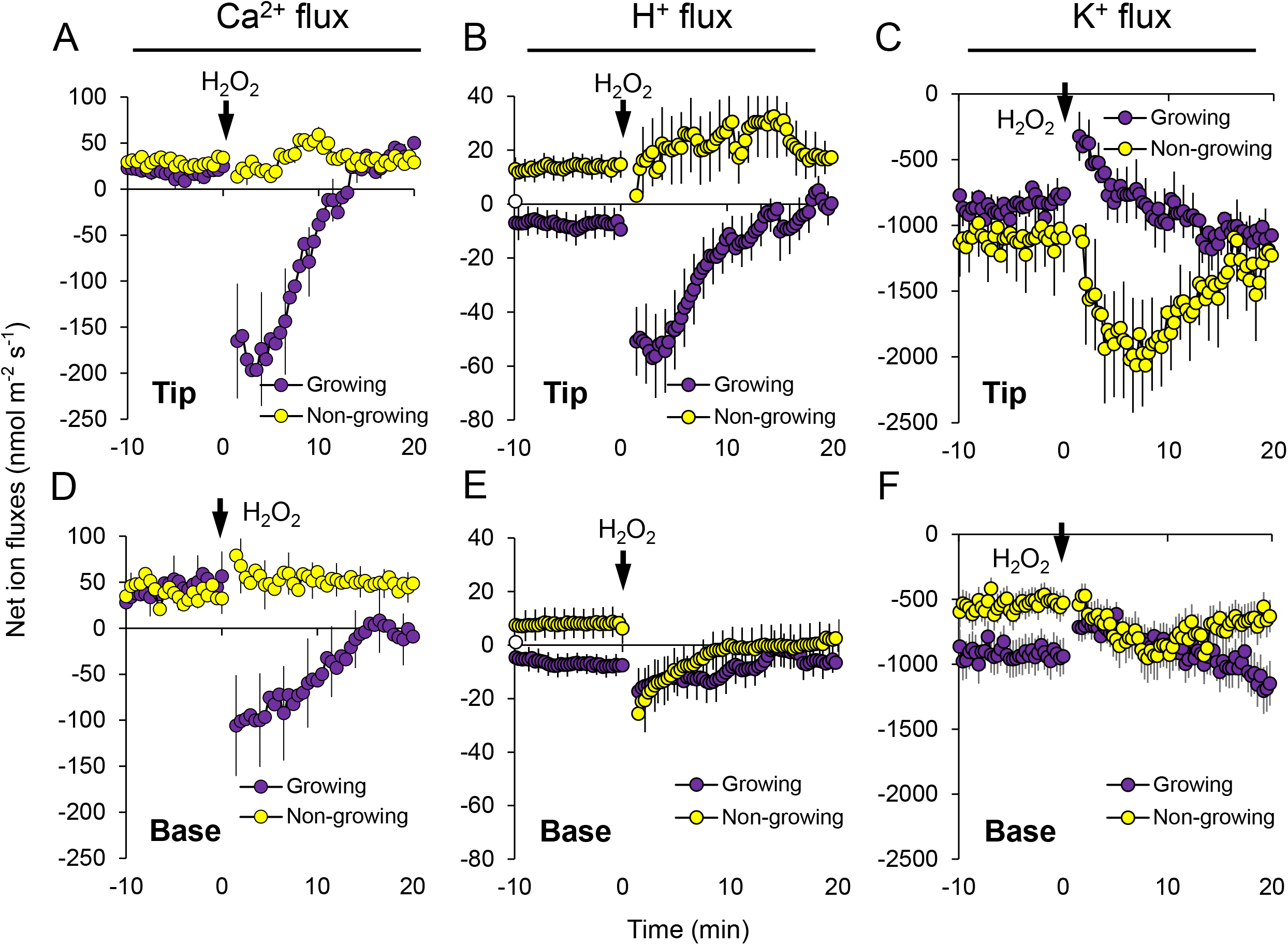
Transient responses of net Ca^2+^ (A and D), H^+^ (B and E) and K^+^ (C and F) fluxes to 5 mM H_2_O_2_ treatment measured from growing (10 DPA) and non-growing (20 DPA) fiber cells in tip and base regions. Error bars indicate SE (n = 5 to 8). The sign convention is “influx positive”.

In the growing fibers, H_2_O_2_ treatment stimulated massive net efflux of Ca^2+^ (Fig. 5, A) and H^+^ (Fig. 5, B), indicative of stimulation of respected ion pumps and resulted in a significant shift towards net K^+^ uptake (less efflux; Fig. 5, C) that could be essential for both turgor-driven growth and a charge balance. No significant effects of H_2_O_2_ on Ca^2+^ and H^+^ fluxes were observed in the non-growing cells, and H_2_O_2_ treatment here resulted in increased net K^+^ efflux from the cell. Thus, it appears that desensitization of the cell and a loss of its ability to respond to H_2_O_2_ may be causally related to the fiber growth mechanism. The magnitude of H_2_O_2_ -induced Ca^2+^ flux response was ~2 fold higher in the fiber tip than that in the base (Fig. 5, A and D) in growing cells. Similar patterns were also observed in the magnitude of H_2_O_2_ -induced changes in net H^+^ flux (Fig. 5, B and E). Here, growing cells showed intrinsically more negative H^+^ flux values (net H^+^ efflux) compared with non-growing cells (net H^+^ uptake). Application of H_2_O_2_ has further increased net H^+^ efflux in growing cells, with much stronger effects reported for the tip (Fig. 5). These findings are consistent with the above observations of higher H^+^-ATPase pumping activity in the tip region, once again pointing at likely higher metabolic activity in this zone. Net K^+^ responses to H_2_O_2_ in the base were qualitatively similar to those in the tip, although several fold lower in magnitude (Fig. 5, C and F).

### Cotton fiber elongation is dependent on Ca^2+^ channel activities

Ca^2+^ flux and its cytosolic homeostasis are largely dependent on the orchestrated activities of numerous plasma membrane Ca^2+^ channels and efflux systems (Ca^2+^_ATases and CAX Ca^2+^/H^+^ exchangers; Bose et al., 2011; Demidchik et al., 2018). To assess a possible role of these transporters in cotton fiber elongation, cotton seeds at 0 or 6 DPA were transferred into the BT medium containing one of the following three blockers-ruthenium red (RR; a known blocker of the tonoplast Ca^2+^-permeable slow vacuolar channels; Pottosin et al., 1999), verapamil (VP; a blocker of the plasma membrane-based voltage-dependent Ca^2+^ channel; De Vriese et al., 2018) and gadolinium (Gd^3+^; a blocker of non-selective cation permeable NSCC channels; Demidchik and Maathuis, 2007; Zepeda-Jazo et al., 2011). The first batch of seeds were cultured from 0 to 6 DPA to examine the impact of the blockers on early fiber elongation. As shown in Figure 6 (A), application of RR and VP reduced fiber elongation by about 20% in comparison with the control with no effect on seed fresh weight, whereas inclusion of Gd^3+^ blocked fiber growth completely and severely inhibited seed growth (Supplemental Figure 1). We further tested the effect of these Ca^2+^ channel blockers on mid stage fiber elongation, namely from 6 DPA onwards. To this end, application RR, VP and Gd^3+^ reduced fiber elongation by about 40%, 75% and 60%, respectively with moderate inhibitory effect on seed weight by RR or VP, but not by Gd^3+^ treatment (Fig. 6, B; Supplemental Figure 1). These findings indicate the essential roles of Ca^2+^ channels in the cotton fiber elongation.

**Figure 6.**
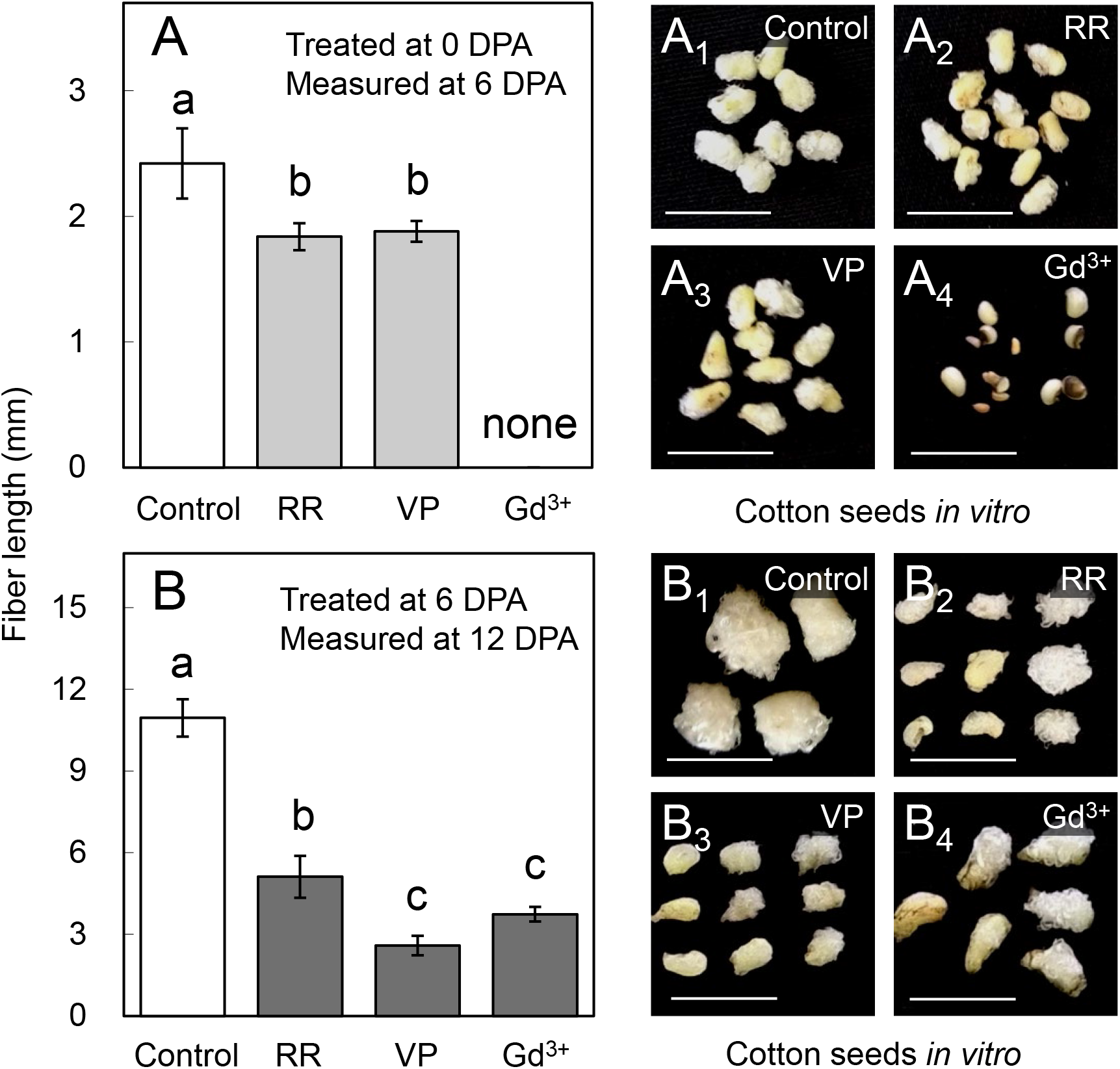
Effects of Ca^2+^ channel blockers on *in vitro* cotton fiber growth. Cotton seeds at 0 or 6 DPA were cultured in the BT medium contains either 50 μM Ruthenium Red (RR), or 200 μM Verapamil (VP), or 1 mM Gadolinium (Gd^3+^) for 6 d. Data labelled with different low-case letters is significant at *P* < 0.05 (One-way ANOVA, Duncan’s test). Error bars indicate SE (n=10-20).

## Discussion

Plant cells grow diffusively across the whole cell surface (Geitmann and Ortega, 2009; Tian et al., 2015) or specifically at the tip area only (Rounds and Bezanilla, 2013; De Jong et al., 2019). The former is characterized with incorporation of a new wall material uniformly distributed across the cell surface leading to even growth over the cell axis as observed in most cell types including, for example, leaf mesophyll cells, epidermal trichomes and pavement cells (Smith and Oppenheimer, 2005; Yanagisawa et al., 2015). Some plant cells concentrate their wall extension through incorporating new wall material only to the apical site, hence becoming tip-growth such as that in the pollen tubes (Qu et al., 2012; Gu et al., 2015) and root hairs (Monshausen et al., 2008; Fan et al., 2011). As outlined in the Introduction, it remains unresolved as whether cotton fiber cells, the most important cell types producing cellulose for the gloable textile industry, follow a tip- or diffusive-based growth pattern. Here, we provided compelling cell biology and eletrophysiology evidence that elongating cotton fiber employs a diffusive but not tip growth mechanism as schematically illustrated (Fig 7).

**Figure 7.**
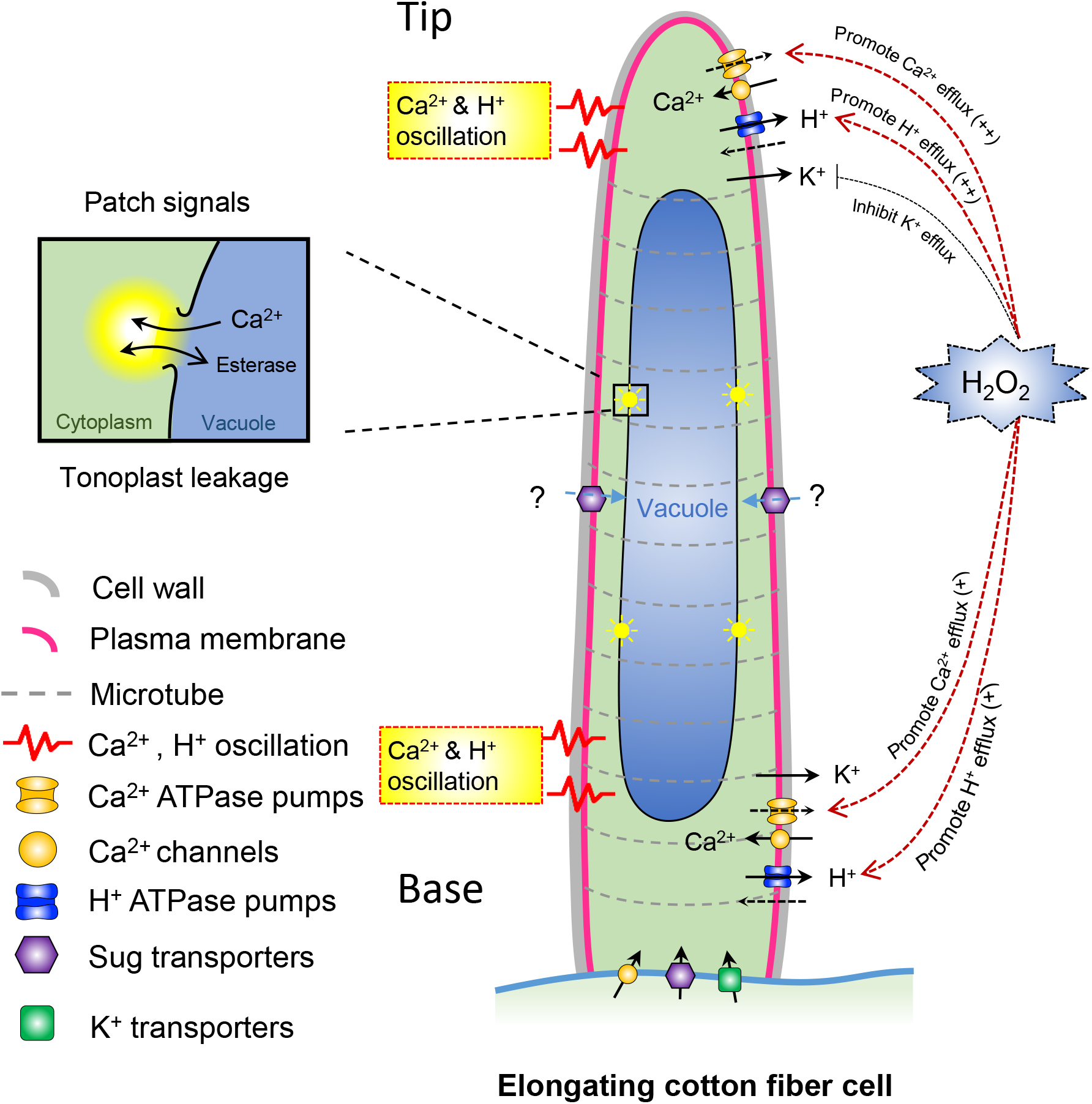
A schematic model on diffusive growth of cotton fiber cells characterized by Ca^2+^ patterning and ion fluxes. Cytosolic Ca^2+^ was found to be evenly distributed from tip to basal regions with Ca^2+^ and H^+^ oscillations detected in both areas in the elongating fiber cells, which also responded to H_2_O_2_ treatment by a massive increase in the net Ca^2+^ and H^+^ efflux in both tip and basal regions. By contrast, non-growing cells (20 DPA) did not respond to H_2_O_2_ treatment and lacked Ca^2+^ and H^+^ oscillations (see text for more details). The findings indicate that the desensitization of the fiber cell and a loss of its ability to respond to H_2_O_2_ may be causally related to the termination of the cotton fiber elongation. Basal location of sugar and k^+^ transporters was based on Ruan et al (2001).

### Cotton fibers follow a diffusive growth as evidenced by a uniform distribtion of the intracellular free cytosolic Ca^2+^ and similar Ca^2+^ flux patterns in both tip and basal zones

By confocal imaging of the Ca^2+^-bindig fluorescent dye, Fluo-3, we showed that Ca^2+^ was uniformely distributed in the cytosol of the fiber cells throughout its elongation period from 0 to 20 DPA (Figs. 1 to 3). Occusionally, less than 5% of young fiber cells appeared to show tip-localized Ca^2+^ at 2-5 DPA, which probably was an artfact derived from the the cytosolic Ca^2+^ being pushed to the tip area during sectioning and imaging process. It is intersting to note from 10 DPA onwards, an increased proportion of the fiber cells exhibited patchy fluorescence of Ca^2+^-binding Fluo-3, which reached ~ 80% by 15 to 20 DPA (Fig. 3, E), when fiber cells enter the transition from elongation to cell wall cellulose synthesis (Ruan, 2007). The fluorescence of Fluo-3 is dependent on esterase activity which catalyzes the removal of the ester group of the non-fluorescent dye Fluo-3 ester once it has diffused from the cell wall matrix to the cytosplasm, thereby allowing Fluo-3 to bind Ca^2+^ leading to emmission of fluorescence (Zhang et al., 1998). Previous studies indicate that as cotton fiber cells approach to the end of the elongation at ~ 15 DPA, the tonoplasts become increasingly leaky, which may generate highly localized pH chnage and even release additonal esterase from the vacuole (Ruan et al., 2001; Ruan et al., 2004), resulting in concentrated Fluo-3 in those areas, hence patchy fluorescence (Fig. 3).

In contrast to tip-based growing cells where Ca^2+^ oscillations are restriced to the apical region, the growing cotton fiber cells displayed Ca^2+^ and H^+^ ossiliation in both tip and basal regions at 5 and 10 DPA but not at 20 DPA when elongation has completed (Fig. 4, Fig. 7). Ion flux oscilaltions are firmly associated with a broad range of developmental and adaptive processes, and rapid (ultradian) oscillations are considered as a hallmark of each elongating cells. Indeed, a strong association was found between oscillations in H^+^ and Ca^2+^ fluxes into epidermal cells and root growth rate (Shabala et al., 1997; Shabala and Newman, 1997), with no oscillations found in roots growing slower than 2 μm min^−1^. These oscillations were always observed in the elongation and meristematic regions of plant roots, but only occasionally in the mature root zone (Shabala and Knowles, 2002). The same is true for the polarized cell growth of pollen tube and root hair in which tip-focused Ca^2+^ oscillations specify the signalling events for rapid cell elongation (Tian et al., 2020).

A critical component of ion flux oscilaltions are H^+^-ATPase pumps that energize membranes and form a feedback loop(s) with various voltage-dependent cation and anion channels (Hansen, 1978; Gradmann, 2001; Shabala et al., 2006). By controlling voltage-gated Ca^2+^ channels (Demidchik et al., 2018), such oscillations in the H^+^-pump activity may rapidly modulate cytosolic free Ca^2+^ concnetrations in the cell creating specific Ca^2+^ “signatures” and activating an array of signalling pathways (Tian et al., 2020). Here we showed that, contrary to reports for pollen tubes or root hairs, such Ca^2+^ and H^+^ flux oscillations were observed in both tip and basal regions of the growing cotton fiber cell (Fig. 4, D) but only in growing cells (Fig. 4, E). The disturbance to cell’s ability to transport Ca^2+^ across the plasma membrane (by pharmacological agents) resulted in a significant (up to 70%) reduction in the fiber growth (Fig. 6). Taken together, this data indicates that elongating cotton fiber employs a diffusive, but not tip growth, mechanism that is strongly dependent on external Ca^2+^ transport from the apoplast across the plasma membrane. It also appears that oscilations in the fibre tip are faster than in the base, suggesting a possibility of the frequency encoding of growth-related signals. The details of this process warrants a separate investigation.

### Growing and non-growing cotton fibers exhibit different flux dynamics of Ca^2+^ H^+^ and K^+^ and their repsonse to H_2_O_2_

H_2_O_2_ has been shown to function as a signalling molecule to promote cotton fiber elongation, likely through enhancing the activity of Ca^2+^-binding protein, Calmodulin (Tang et al., 2014). H_2_O_2_ is also known as a potent activator of several types of Ca^2+^-permeable ion channels, forming so called ‘Ca^2+^-ROS hub” and amplifying cytosolic Ca^2+^ signals (Demidchik et al., 2018) in root epidermis and leaf mesophyll cells. Here, however, application of H_2_O_2_ resulted in a massive net Ca^2+^ efflux (Fig. 5, A and D) that was observed in both apical and basal region but only in growing fibers. As electrochemical [Ca^2+^] gradient across the plasma membrane favours thermodynamically passive Ca^2+^ uptake, the observed net Ca^2+^ efflux can be only a result of operation of some active Ca^2+^ transport system (such as Ca^2+^-ATPase or Ca^2+^/H^+^ exchanger) activated by H_2_O_2_. Zepeda-Jaso et al. (2011) reported that hydroxyl radicals (another type of ROS) caused activation of eosin yellow-sensitive (a specific Ca^2+^ pump inhibitor) Ca^2+^ efflux from pea root epidermis, and that these effects were potentiated by polyamines (Velarde-Buendía et al., 2012). Thus, we speculate that a similar scenario may be applicable here.

The physiological rationale behind activation of Ca^2+^ efflux remains a subject of a separate study. ACA-type (plasma membrane-based) Ca^2+^-ATPases are essential component of the polarized cell growth. Four ACAs (ACA2, 7, 9, 11) are highly expressed during most of the pollen developmental stages (García Bossi et al., 2020), and insertional mutants of ACA9 show reduced pollen tube growth (Schiøtt et al., 2004). Ca^2+^-ATPases are also essential for a root hair growth. In our case, net Ca^2+^ efflux was activated by ROS treatment (Fig. 5). The most likely explanation on the stimulation of Ca^2+^ efflux by H_2_O_2_ is that such stimulation of Ca^2+^ ATPase is required to return cytosolic [Ca^2+^] levels down to the basal levels, once the ROS signalling is over (Bose et al., 2011). Indeed, the above “Ca^2+^-ROS hub”, that is formed by the H_2_O_2_-inducable Ca^2+^ permeable non-selective cation channel and NADPH oxidase (Demidchik et al., 2018), operates in a positive feedback manner and can result in an uncontrollable increase in ROS accumulation in the growing cell, with a danger of a possible damage to key macromolecules and cellular structures. Hence, once the ROS signalling is over, operation of this self-amplifying system should be ceased. The H_2_O_2_-induced activation of Ca^2+^-ATPase may serve just this purpose.

The elongating (5 and 10 DPA), but not elongated (20 DPA), cotton fibers exhibited net H^+^ efflux, with the strongest efflux observed at 10 DPA (Fig. 4). This efflux was vanadate-sensitive (data not shown), implying involvement of H^+^-ATPase. The magnitude of H^+^ efflux was further stimulated by H_2_O_2_ treatment (Fig. 5) implying activation of H^+^-ATPase (in addition to that of Ca^2+^ ATPase). This activation would hyperpolarize the plasma membrane and positively impact on acquisition and transport of essential nutrients and metabolites (Ruan, 2007).

Cotton fiber growth relies on the import of nutrient resources from the basal ends connecting the underlying seed coat. During fiber elongation, the cellular pathway for nutrient import switches from symplasmic pathway via plasmodesmata early in elongation to apoplasmic route at ~10 DPA at the onset of rapid fiber elongation (Ruan et al., 2001; Ruan et al., 2004). This switch coincides with strong expression of a group of plasma membrane sugar and K^+^ transporters to uptake sugars and K^+^, major osmotical solutes in the fiber cells (Ruan et al., 2001; Zhang et al., 2017b). Here, we show that H_2_O_2_ treatment resulted in a shift towards net K^+^ uptake in a growing cotton fiber (Fig. 5, C and F) that could be explained by both H^+^-ATPase pump-mediated membrane hyperpolarization (hence, opening inward-rectifying K^+^ channels) and increased pH gradient across the plasma membrane to provide a driving force for operation of the high affinity K^+^ uptake systems (e.g. K^+^/H^+^ exchangers; Rubio et al. 2020). The highest H^+^-ATPase activity and its increased sensitivity to H_2_O_2_ at 10 DPA may be also essential to transport sucrose (via H^+^/sucrose symporter (Ruan et al., 2001; Zhang et al., 2017b). Together with K^+^, sucrose import may also osmotically drive fiber elongation as K^+^ and malate account for ~ 50% of the total osmolality with soluble sugars making up the remaining 50% and they collectively play essential role in generating osmotic potential, hence turgor (Dhindsa et al., 1975; Ruan et al., 2001; Wang et al., 2010). In this context, activation of H^+^-ATPase by ROS could be a critical step in generating cell turgor for fiber elongation.

## Materials and Methods

### Plant Material

Cotton (*Gossypium hirsutum* cv. Coker 315 plants were grown under controlled conditions as previously described (Ruan et al., 1997). Cotton seeds were sown in a potting mixture (Metro-Mix 200 growing medium, Scotts, Columbus, OH). The plants were raised under greenhouse conditions with partial temperature control (25-30 °C during the day for 14 h and 18-22 °C during the night for 10 h. About 100 g per pot of Osmocote (Scotts), a controlled release fertilizer with N:P:K at 1:1:1, was applied once every 20 d. The plants were watered once every 2 d. Cotton bolls were sampled at 9:00~ 10 am on d of anthesis for ovule culture.

### Cotton Ovule Culture

The procedure of cotton ovule culture was carried out as described (Li et al., 2010) with some modifications. Cotton bolls (0 DPA) at 1^st^ nodes of middle fruiting branches were collected and surface-sterilized with 70 % (v/v) ethanol for 30-60 s and a quick exposure to methane burner flame, followed with treatment of 6% (v/v) sodium hypochlorite (NaOCl) for 20 min. Thereafter, the bolls were washed with sterile water to remove residual NaOCl. Ovules were removed aseptically from middle part of each locule and floated on 50 ml liquid BT medium (Beasley and Ting, 1974) in a 300 ml- plastic culture bottle with 20 ovules per flask. The bottles were placed in the dark at 30 °C without shaking until observations.

The BT culture medium was prepared according to (Beasley and Ting, 1974). indole-3-acetic acid (IAA) sodium salt and gibberellic acid (GA_3_) potassium salt were dissolved in water and filter sterilized to obtain stock solutions of 50 mM and 5 mM, respectively. In practice, 5 μM IAA and 0.5 μM GA_3_ were added to the BT medium after autoclaving. All above operations were performed under sterile condition.

### Confocal imaging of Ca^2+^ patterning

Cotton ovules with fibers attached were pre-loaded with the intracellular Ca^2+^ sensitive fluorescent probe, fluo-3 acetoxymethyl (Fluo-3 thereafter) ester (Biotium, US) following a protocol adapted from (Zhang et al., 1998). For loading, cotton ovules were incubated in the BT medium contains 20 μM Fluo-3 ester (stock in DMSO), 200 μM CaCl_2_, and 50 mM sorbitol in a 12-well cell culture plate (2 ml per well) for 3 h at 4 °C in the dark to minimize the hydrolysis of AM ester by potential extracellular esterases. Cotton ovules were then transferred to a new BT medium incubated for 2 h at 26 °C to allow the cleavage of the loaded Fluo-3 ester to Fluo-3 by cytosolic esterases, thereby trapping the impermeable Fluo-3 in the cytosol of fiber cells. Afterwards, hand-cut sections of cotton ovule with fiber attached were washed 3 times by phosphate-buffered saline (PBS) plus 100 mM sucrose. In specified instances, sections were counterstained with 0.1 % (w/v) Calcofluor White for 30 s to label the cell wall and the (Zhang et al., 2015). Thereafter, sections were transferred to 1 ml of PBS containing 100 mM sucrose for preparation of confocal imaging. Sections of cotton ovules were carefully placed on slides to ensure most fiber cells were placed on the same focal plane.

Fluorescent imaging of cotton ovule and fiber cells was performed using an Olympus FV1000 confocal laser scanning microscopy (Olympus, Japan). Fluo-3 was excited with a 488 nm diode laser (15 mW, laser power set to 50 %) and emitted fluorescence captured at 515 nm, while Calcofluor White was excited with a 405 nm UV laser (50 mW, laser power set to 15 %) and emitted fluorescence captured at 460 nm. Gain of the photomultiplier tube was set to 600 V for Fluo-3, and 400 V for Calcofluor White.

To imaging Fluo-3 fluorescence in cotton root hairs, 14 d old seedlings were chosen for root sampling, and loading of Fluo-3 as that for cotton ovules. For imaging of Fluo-3 in pollen tubes, pollen grains were harvested from blooming flowers and suspended in the culture medium (1.0 mM KCl, 0.06 mM CaCl2, 1.0 mM H3BO3, and 1.0 mM MES, pH 6.0 (Tris)) at room temperature (26 °C) for 30 min. Then Fluo-3 ester (20 μM final concentration) was added to the culture medium and incubated for 2 h at 4 °C in darkness. Pollen grains were then washed three times with dye-free culture medium and then incubated for 2 h at 26 °C before fluorescent imaging.

### Plasmolysis of fiber cell

To better identify the subcellular localization of the Ca^2+^ signal, Fluo-3 loaded cotton ovules (5 DPA) were counterstained with 10 μM RH-414, a cell membrane tracker (Molecular Probes, US), during the last 30 min of cleaving Fluo-3 ester, before hand-cutting. Thereafter, sections of cotton ovule were soaked in the PBS solution containing 500 mM or 1 M sorbitol for 20 min to achieve moderate or severe plasmolysis of thefiber cells, respectively. RH-414 was excited with a 559nm diode laser (15 mW, laser power set to 25%) and emitted fluorescence captured at 625 nm. Gain of the photomultiplier tube was set to 500 V.

### Non-invasive H^+^, Ca^2+^ and K^+^ flux measurements using MIFE^TM^ technique

The H^+^, Ca^2+^ and K^+^-selective microelectrodes were prepared and calibrated as described elsewhere (Shabala et al., 2013). A lock from the bolls of 5 DPA or 10 DPA or 20 DPA was gently removed and secured on a glass slide using a thin paraffin strip and conditioned in basal salt medium (BSM; 120 mM Glucose, 1 mM KCl, 0.1 mM CaCl_2_ pH 5.7 ± 0.2) for 40 minutes at room temperature for recovery of growth (Supplemental Figure S1). The net H^+^, Ca^2+^ and K^+^ fluxes were measured from apical and basal regions of the cotton fiber at 40 μm away from individual fiber cells using MIFE technique for up to 60 min. To study the H_2_O_2_-induced flux changes, steady-state ion fluxes were measured for 10 minutes at respective regions of the cotton fiber. Thereafter, 5 mM H_2_O_2_ was added to the BSM, and transient ion fluxes were monitored for further 30-40 min.

### Pharmacological experiments

To examine the role of Ca^2+^ channels in cotton fiber growth, the pharmacological approach was undertaken. Ruthenium red (RR; known blocker of the tonoplast Ca2+-permeable slow vacuolar channels), verapamil (VP; a blocker of the plasma membrane-based voltage-dependent Ca^2+^ channel) and gadolinium (Gd^3+^; a blocker of non-selective cation permeable NSCC channels were added to the culture medium individually at specified DPA. RR and Gd^3+^ were added from aqueous stocks of 5 mM and 50 mM, respectively; VP was added from a 100 mM stock in ethanol. The pharmaceuticals were diluted prior to use to working concentrations of 50 μM, 0.2 mM, and 1 mM for RR, VP, and Gd^3+^, respectively.

### Measurement of the fiber length

Cotton ovules with fibers attached were boiled in 100 ml of distilled water with 2-3 drops of 1 M HCl inside for 1 min. Afterwards, each ovule was carefully placed on a convex dish, and fibers grown on the ovule were streamed out with a jet of water. Then, fiber length was determined from the attachment point of fiber and ovule to the edge where most fibers terminate at. A vernier caliper together with a dissecting microscope were employed for the measurement.

### Data analysis

Fluorescent images captured by confocal microscope were edited in software FV10-ASW 4.0 viewer. Schematic diagram was drawn by software PowerPoint 2013 (Microsoft). Graphs were plotted by Excel 2013 (Microsoft). Student’s T test and ANOVA analyses were performed in SPSS 20 (IBM).

## ACKNOWLEDGMENTS

J-S Y gratefully acknowledges support from China Scholarship Council. We thank Dr Huiming Zhang for advice on confocal imaging of Ca^2+^ patterning.

## Supplemental Data

**Supplemental Figure S1.** The effect of Ca^2+^ blockers on cotton ovule growth.

